# How do Cockroach Groups Integrate Multiple Attributes in a Best-of-N Task?

**DOI:** 10.64898/2026.07.24.740418

**Authors:** Paco C.K. Chow, Horacio G. Rotstein, Simon Garnier

## Abstract

Collective decision-making experiments have largely focused on simple scenarios in which groups choose between two options that differ along a single attribute. However, it remains unclear whether principles derived from this single-attribute best-of-2 paradigm generalizes to the multi-attribute, multi-option decisions that groups often face in nature. Here, we use mean-field and agent-based models of cockroach aggregation to study collective decision-making across a range of problem complexities, varying both the number of options (*N*) and the number of attributes describing each option. Our models make 3 novel predictions: (1) cockroach groups use a compensatory algorithm when integrating attributes, trading off strength in one attribute against weakness in another; (2) decision-making collapses abruptly once *N* exceeds a critical threshold; and (3) decision time scales non-monotonically with *N*. Together, these results indicate that dynamics characterized in single-attribute best-of-2 experiments do not extrapolate to multi-attribute best-of-*N* decision-making. Collective choice under realistic complexity may follow principles not yet captured by existing models.

## 1 Introduction

Animal groups must constantly make collective decisions on where to forage, migrate, and nest (Sumpter, 2006; Garnier and Moussaïd, 2022; Sasaki and Pratt, 2018). For example, cockroach groups search for the darkest shelter to aggregate under (Canonge et al., 2011; Freeberg and Fiset, 2023); ant colonies congregate at high-quality foraging sites (Beckers et al., 1990; Kolay et al., 2020); honeybee swarms look for new nest sites to build homes together (Franks et al., 2002; Seeley, 2010). These types of decisions are known as the best-of-*N* problem (Valentini et al., 2017), where a group must choose the most suitable option from a set of *N* alternatives.

Solving best-of-*N* problems in the wild often involves several—sometimes conflicting—attributes (e.g., food quality and distance from nest) that the group must integrate to make the best possible choice. In addition, animal groups must typically be able to do this across a wide range of options. While there have been countless empirical studies on collective decision-making in animal groups, these studies have largely focused on simple search problems, typically involving only two options that differ in a single attribute (Latty and Trueblood, 2020; Kubo et al., 2022; Visscher, 2007; Jeanson et al., 2012). Our current understanding of collective decision-making is, therefore, largely based on an extremely restricted region of the decision landscape that may not capture how animal groups operate in the real world.

Here, we define **decision complexity** as the number of elements that need to be considered to solve a best-of-*N* problem. Decision complexity can be increased in two ways: (1) by increasing the number of available options, and (2) by increasing the number of differing attributes that need to be factored into the decision-making process. Figure 1 summarizes this in a two-dimensional decision landscape using the example of cockroaches seeking a target shelter among distractors.

**Figure 1:**
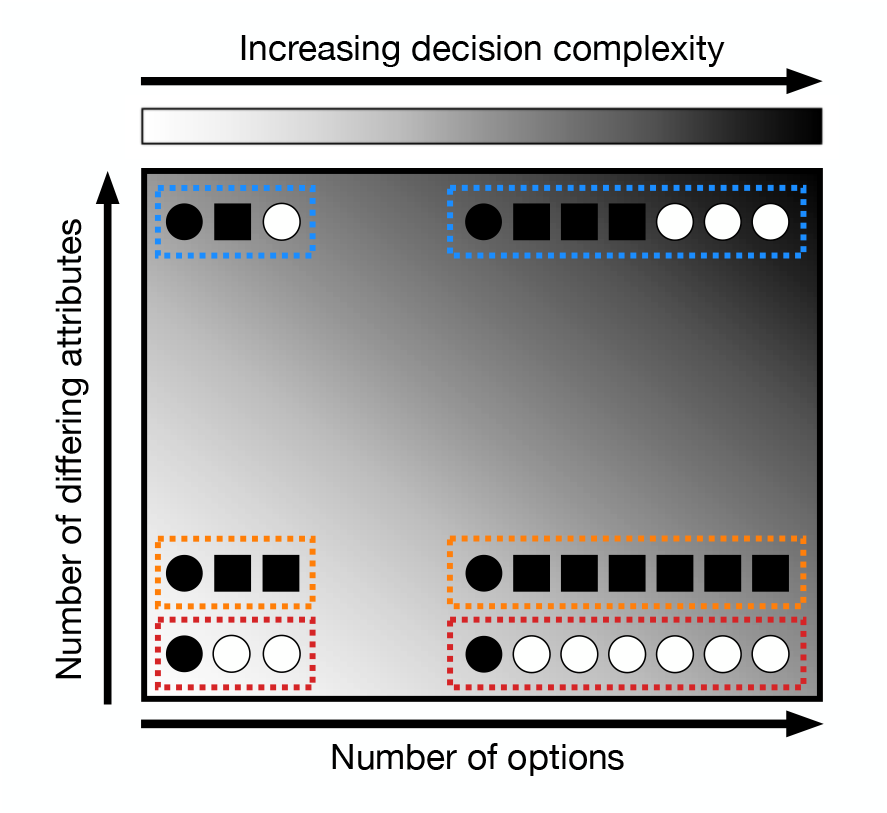
Schematic showing decision complexity of different distractor configurations. Black circles represent the target shelter. Black squares represents large distractors. White circles represent bright distractors. The complexity of the decision problem increases (i) along the *x* -axis, by increasing the number of distractors, and (ii) along the *y* -axis, by increasing the number of differing attributes to be considered simultaneously (e.g., size and brightness). Note that the all large distractors configuration and all bright distractors configuration have the same decision complexity since the number of differing attributes in both cases is 1.

Exploring this complex decision landscape is important to understanding how animal groups handle decision-making problems that are ecologically realistic. More importantly, we can use this to make inferences about the properties of the biological computational algorithms underlying information processing during decision-making. For example, by presenting different options with attributes that require computing trade-offs to find the best option, one can infer whether the group is using an algorithm that allows an option that is less attractive in one key attribute to compensate by being more attractive in other less important attributes (Franks et al., 2003). In addition, by increasing the number of options and measuring the decision time, one can infer the extent with which the group is processing attribute information across options in parallel (as opposed to in series) (Treisman and Gelade, 1980).

Here, we use a classical model of collective decision-making in cockroach groups that had been studied in single-attribute binary-choice experimental setups (Ame et al., 2004; Amé et al., 2006; Campo et al., 2011). We show that exploring decision complexity within this framework results in counter-intuitive predictions at the level of the group.

Cockroaches are social insects that aggregate under shelters to reduce the risk of predation, reduce physical stresses such as desiccation, and to accelerate development (Ledoux, 1945; Izutsu et al., 1970; Wileyto et al., 1984). When presented with multiple shelters that vary in brightness and capacity, cockroach groups are able to collectively select and aggregate under the darkest shelter (to minimize risk of predation) that is small enough to fit the group (this maximizes the density of cockroaches under the shelter to reduce desiccation) (Canonge et al., 2009; Jeanson and Deneubourg, 2007). Existing work has shown that this process is an entirely self-organized process: individuals integrate social and environmental cues through local interactions (Conradt and Roper, 2005; Jeanson et al., 2005), allowing the group to achieve consensus without the need for central control. Specifically, the probability that a cockroach leaves a shelter depends on both social (decreases with increasing density of individuals under that shelter) and environmental (increases with increasing brightness) factors (Amé et al., 2006).

These cockroach-cockroach and cockroach-environment interactions have been separately modeled to analyze the results of best-of-2 experiments (Amé et al., 2006; Campo et al., 2011; Canonge et al., 2009). Here, we incorporated both types of interactions into a single mean-field model that is able to integrate information about both shelter density and brightness across *N* options. We used this model to (1) identify which attribute the cockroaches value more, (2) determine whether cockroaches are able to compute trade-offs in attribute attractiveness, and (3) understand how the decision-making dynamics of the group scale with best-of-*N* problems of increasing numbers of options. Our results show a non-trivial relationship between the number of options available and the decision time.

## 2 Methods

### 2.1 Mean-field model

A population of *P* cockroaches choose among *N* shelters that vary in shelter capacity (*ϕ*) and brightness (*θ*). Let *x*_*i*_ be the number of cockroaches under shelter *i*, and *X*_*i*_ be the proportion of the cockroach group under a shelter 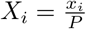. The shelter capacity *ϕ* is defined as a scaling factor relative to the group size *P* . A shelter *i* that is just small enough to fit the group has *ϕ*_*i*_ = 1, and for larger shelters *ϕ*_*i*_ *>* 1. Similarly, the darkest shelter has base brightness *θ*_*i*_ = 1, and for brighter shelters *θ*_*i*_ *>* 1. All cockroaches begin in an uncommitted state.

The dynamics of *X*_*i*_ are described by:

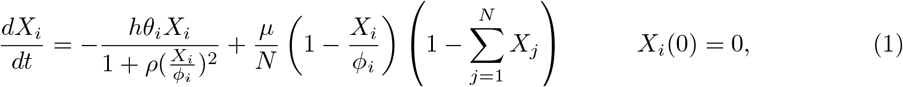

for *i* = 1, … *N, h, ρ* and *µ* are fixed model parameters that govern the sensitivity of cockroaches to light, density and determine the speed at which cockroaches discover new shelters, respectively. As only best-of-2 experiments have been done with cockroaches, we use *h, ρ* and *µ* values that have been fit to existing data from best-of-2 experiments (Amé et al., 2006) to capture the biological timescale of cockroach decision-making dynamics: *h* = 0.01, *ρ* = 1667 and *µ* = 0.002.

In this model, the rate of cockroaches leaving a shelter (the first term) is dependent on the brightness *θ*_*i*_ and the density of cockroaches 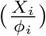 under that shelter, where shelters with low brightness and high density have the lowest leaving rate. The recruitment rate to a shelter (the second term) is dependent on the number of shelters 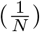, the level of saturation within the shelter 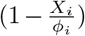, and the number of uncommitted cockroaches 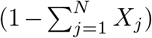. This rate is reduced when there are many available shelters, when the shelter is already full, and when there are few uncommitted cockroaches remaining. Simulations were run until the system converges to a state of equilibrium.

#### 2.1.1 Experiment 1

We first performed a best-of-2 task to examine how cockroach groups process information when there are conflicting attributes. Shelter 1 is attractive in its capacity (*ϕ*_1_ = 1) but unattractive in its brightness (*θ*_1_ *>* 1), and shelter 2 is unattractive in its capacity (*ϕ*_2_ *>* 1) but attractive in its brightness (*θ*_2_ = 1). We systematically varied *θ*_1_ and *ϕ*_2_ from 1 to 12 with intervals of 0.1 to test the preference of the group at different combinations of attribute values. To quantify the preference of the group, we measured the difference in the proportion of the group under shelter 1 and shelter 2 (*X*_1_ − *X*_2_) at the end of the simulation.

#### 2.1.2 Experiment 2

We then simulated another best-of-2 task to examine whether cockroach groups use a compensatory or non-compensatory algorithm when integrating attribute information. Compensatory algorithms allow trade-offs between criteria, meaning that an option that is less attractive in a desirable attribute can still rank highly if it is attractive in other less desirable attributes. Non-compensatory algorithms do not allow trade-offs, so an unattractive attribute alone can lead to the elimination of an option.

Shelter 1 is attractive in its capacity (*ϕ*_1_ = 1) but unattractive in its brightness *θ*_1_ = 2, and shelter 2 is relatively unattractive in its capacity *ϕ*_2_ *>* 1 but relatively more attractive in its brightness 1 *< theta*_2_ *<* 2. We systematically varied *ϕ*_2_ and *θ*_2_ and measured the difference in the proportion of the group under shelter 1 and shelter 2 (*X*_1_ − *X*_2_) at the end of the simulation.

#### 2.1.3 Experiment 3

For experiment 3, we simulated best-of-*N* tasks with 1 target and *N* − 1 distractors. In our protocols, shelter 1 is the target (*X*_1_ = *X*_*T*_) and the remaining shelters are the distractors. For future use, we define the proportion under all distractors with 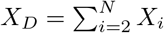. The target shelter is attractive in both its capacity (*ϕ*_*T*_ = 1) and its brightness (*θ*_*T*_ = 1). In our simulations, all distractors are either unattractive in their capacity or their brightness. In our simulations, distractors unattractive in their capacity (large distractors) all have capacity *ϕ*_*D*_ = 1.75 and brightness *θ*_*D*_ = 1. Distractors unattractive in their brightness (bright distractors) all have capacity *ϕ*_*D*_ = 1 and brightness *θ*_*D*_ = 1.75.

*ϕ*_*D*_ and *θ*_*D*_ control the difficulty of the best-of-*N* problem, where increasing *ϕ*_*D*_ and *θ*_*D*_ decreases the difficulty of finding the target shelter among the distractors as it decreases the contrast between the target and the distractor.

For each best-of-*N* experiment, the set of distractors can be of various types with different decision complexities: Distractors can be either all too large or all too bright (low decision complexity), or half too large but dark and half too bright but the right size (high decision complexity). This is illustrated in Figure 1. The final proportion is defined as the proportion of the group under the target shelter at the end of the simulation. To measure the decision time of the group, we define *T*_95_ as the time taken to reach 95% of the final proportion.

### 2.2 Agent-based model

As the mean-field model is continuous and assumes large group sizes, we implemented an agent-based model (ABM) with the same local interactions. This is a more realistic model that will allow us to validate the results with discrete agents and smaller group sizes.

Similarly to the mean-field model, there are *P* agents and *N* shelters, where each shelter *k* ∈ 1, 2, …, *N* has brightness *θ*_*k*_ and capacity *ϕ*_*k*_. The state at time *t* for an agent *i* is characterized by *A*_*i,t*_ where *A*_*i,t*_ ∈ *{*0, 1, 2, …, *N}*. If *A*_*i,t*_ = 0, the agent is uncommitted. If *A*_*i,t*_ = 1, the agent is under the target shelter. If *A*_*i,t*_ = 2, …, *N*, the agent is under a distractor shelter. We define *X*_*k*_ as the proportion of agents under shelter *k*. We use Δ*t* = 0.1 to accurately capture dynamic interactions.

All the agents start in an uncommitted state *A*_*i*,0_ = 0. At each iteration, uncommitted agents either remain in the uncommitted state or discover a shelter. The probability of transitioning into the shelter scales with the density of individuals under the shelter, meaning that it is harder for an agent to enter a crowded shelter. Thus, if *A*_*i,t*_ = 0:

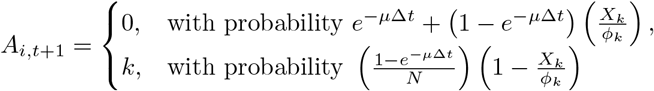

for *k* = 1, … *N*

Alternatively, if an agent is under a shelter *k* (*A*_*i,t*_ = *k*), *k* = 1, …, *N*, we define the leaving rate 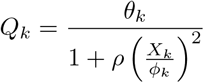, and

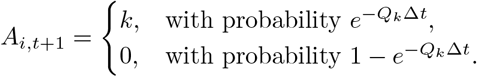

### 2.3 Spatial agent-based model

The ABM implemented thus far is spatially agnostic as each shelter is equally likely to be discovered. We introduce a spatial dependence by using a transition matrix *T* ∈ **R**^*N×N*^ where the probability of transitioning from one shelter to another depends on their pairwise distance. To empirically derive the transition probabilities, we simulate many random walks in a rectangular field with circular shelters arranged in a triangular grid (see Figure 7A) and measure the frequency of transitioning between every pair of shelter.

The spatial ABM is implemented in exactly the same way as the space-agnostic ABM except for the fact that when *A*_*i,t*_ = 0:

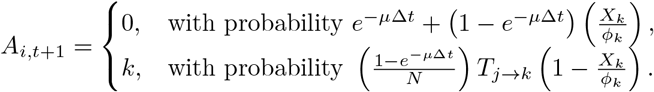

where *T*_*j*→*k*_ is the probability of transitioning from the last shelter *j* the agent was into a new shelter *k*.

## 3 Results

### 3.1 Cockroaches value shelter capacity over brightnesss

How does the model process multi-attribute information about shelters? We first performed an experiment (Experiment 1) that allowed us to identify how the model ranks attributes, i.e. whether the cockroaches value shelter capacity or brightness more. We designed a best-of-2 setup where each option is attractive in one attribute but not the other.

Figure 2A shows a heatmap of the decision made by the group at different combinations of values of *θ*_1_ (brightness of shelter 1) and *ϕ*_2_ (capacity of shelter 2) and the dynamics of decision-making at key regions of the heatmap. We found that cockroaches choose the bright shelters when *θ*_1_ = *ϕ*_2_ i.e. when both shelters have an equally unattractive attribute, up to a threshold. The model therefore predicts that cockroaches value the capacity of the shelter over its brightness, as they are more willing to compromise on a decrease in attractiveness of the brightness attribute than an equivalent decrease in attractiveness of the capacity attribute. Interestingly, as *θ*_1_ and *ϕ*_2_ change, the decision boundary between choosing the shelter 1 and 2 changes non-linearly.

**Figure 2:**
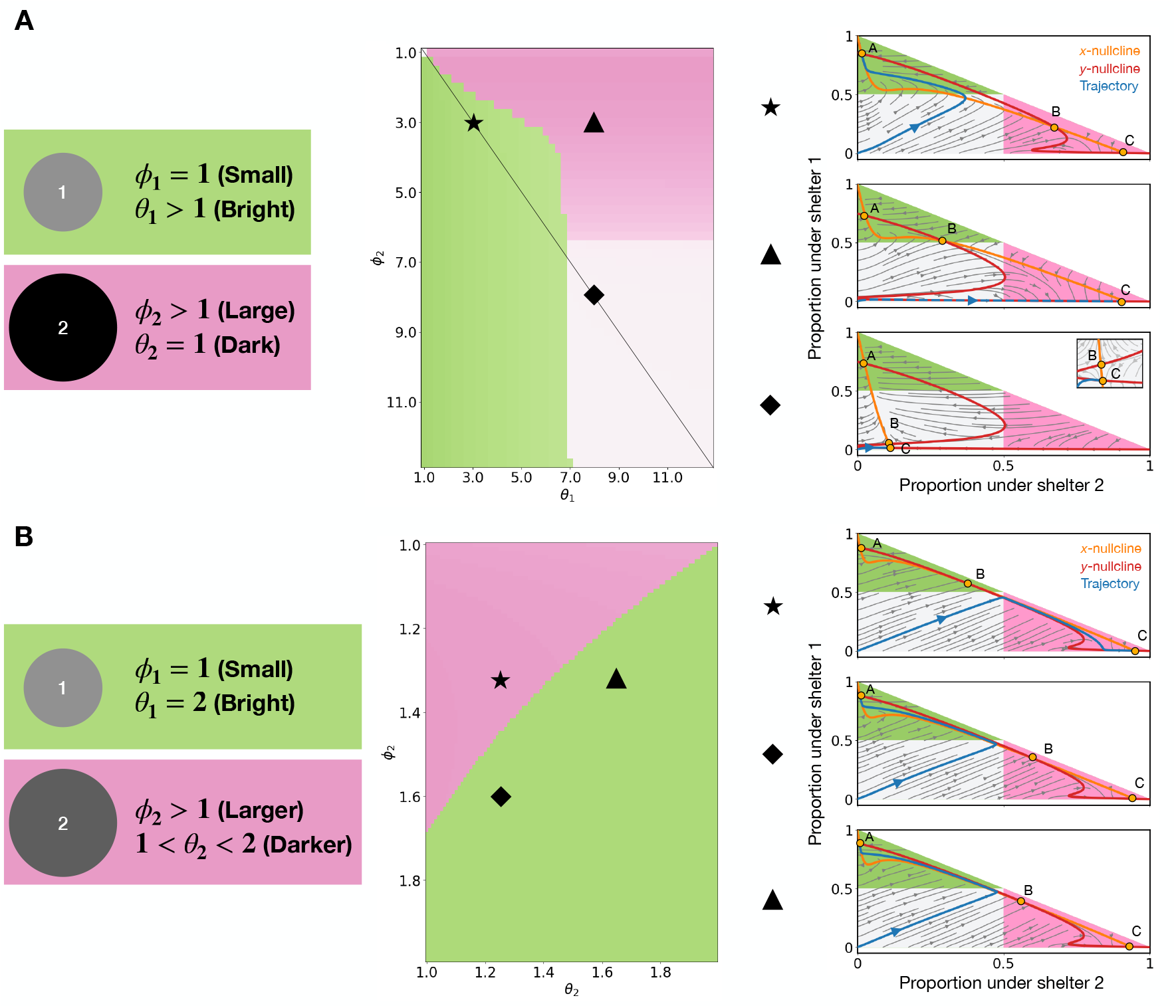
A) (Left) Experimental setup of the best-of-2 experiment. Shelter 1 is attractive in the capacity attribute but unattractive in the brightness attribute. Shelter 2 is unattractive in the capacity attribute but attractive in the brightness attribute. (Middle) Heatmap showing which shelter is picked by the model. Diagonal line shows *ϕ*_2_ = *θ*_2_ . (Right) Phase portraits at corresponding regions. The *x*- and *y*-nullclines are shown in orange and red respectively. Gray region represents the region of indecision. Colored regions represent regions of consensus under shelter 1 (green) or shelter 2 (pink) (i.e., more than half of the group has aggregated under a shelter). Gray arrows represent vector fields and gray curves represent trajectories of the system for different initial conditions. The trajectory when the system starts with all cockroaches uncommitted is shown in blue. B) Equivalent plots to (A) with different best-of-2 experimental setup. Shelter 1 is attractive in the capacity attribute but unattractive in the brightness attribute. Shelter 2 is relatively less attractive in the capacity attribute but relatively more attractive in the brightness attribute. These results have been replicated with our agent-based model (Supplementary Figure 1).

We can understand the dynamics of this by performing a phase portrait analysis at different points on the heatmap (Fig. 2A right panel). The intersections between the two nullclines (zero-level curves) for the variables *X*_1_ and *X*_2_ (proportion of the group under shelters 1 and 2 respectively) define the fixed-points (states of equilibrium). The trajectory converges to one of the different fixed-points depending on the initial condition and the structure of the vector field. As we are only interested in the initial condition where all cockroaches start in the uncommitted state (*X*_1_ = 0, *X*_2_ = 0), we refer to the resulting trajectory as “the trajectory”.

For the range of *ϕ*_1_ and *θ*_2_ values tested, the nullclines intersect at 3 fixed points. Fixed points A and C are stable, while fixed point B is unstable. From the different phase portraits shown, the shape and position of the nullclines depend on *ϕ*_1_ and *θ*_2_. When *ϕ*_1_ = *θ*_2_ = 3 (star), the trajectory moves towards fixed point A, representing a consensus under the bright shelter (shelter 1). As *θ*_2_ increases (from star to triangle), the *y*-nullcline moves to the left. This moves the position of unstable fixed point B to the left, resulting in the trajectory moving towards fixed point C instead, representing a consensus under the large shelter (shelter 2). As *ϕ*_1_ increases (from triangle to diamond), the *x*-nullcline moves downwards. The trajectory still moves towards fixed point C, but its new position represents a state of indecision (since C is found in *X*_1_ *<* 0.5 and *X*_2_ *<* 0.5, i.e., a very low proportion of the group is under shelters 1 and 2).

### 3.2 Cockroaches use a compensatory algorithm to process attributes

Having established how the model ranks attributes, we can infer whether cockroach groups use a compensatory or non-compensatory algorithm when integrating attribute information (Experiment 2) by asking: can a shelter that is less attractive in a highly ranked attribute (the capacity attribute) still be preferred if it is compensated by a sufficient increase in attractiveness of a lower ranked attribute (the brightness attribute)? If cockroaches use a compensatory algorithm, a shelter that is relatively unattractive in the capacity attribute (*ϕ*_2_ *> ϕ*_1_) can still be preferred if it is sufficiently more attractive in the brightness attribute (*θ*_2_ *< θ*_1_). If cockroaches use a non-compensatory algorithm, a shelter that is relatively unattractive in the capacity attribute will never be preferred regardless of the value of the brightness attribute.

From Figure 2B, we found that there are a range of combinations of *ϕ*_2_ and *θ*_2_ values (pink region) such that cockroaches prefer shelter 2 (a shelter that is less attractive in the capacity attribute but more attractive in the brightness attribute) over shelter 1 (a shelter that is attractive in the capacity attribute but less attractive in the brightness attribute), consistent with the predictions of a compensatory algorithm. The phase portraits show that changing *ϕ*_2_ and *θ*_2_ changes the position of unstable fixed point B, which directs the trajectory towards stable fixed points A or C (corresponding to consensus under shelter 1 and shelter 2 respectively).

### 3.3 Distinct processing regimes emerge with increasing distractor number in the mean-field model

#### 3.3.1 Group-level properties

How do cockroaches process multi-attribute information when there are more than 2 shelters? In Experiment 3, we numerically solved the mean-field model with different numbers and types of distractors, and found that cockroaches can collectively decide to aggregate under the target shelter across a wide range of decision complexities. Figure 3A demonstrates that the dynamics of this collective decision-making process differs in a non-monotonic fashion depending on the decision complexity. At low numbers of distractors (top panel), decision-making is relatively slow. Surprisingly, at intermediate numbers of distractors (middle panel), consensus under the target shelter is achieved relatively faster despite the increase in decision complexity. At high numbers (bottom panel) of distractors, the time necessary to reach a collective decision is once again slower. This counterintuitive result is described in more detail in Figure 3B. From this, we observe 3 group-level properties that emerge.

**Figure 3:**
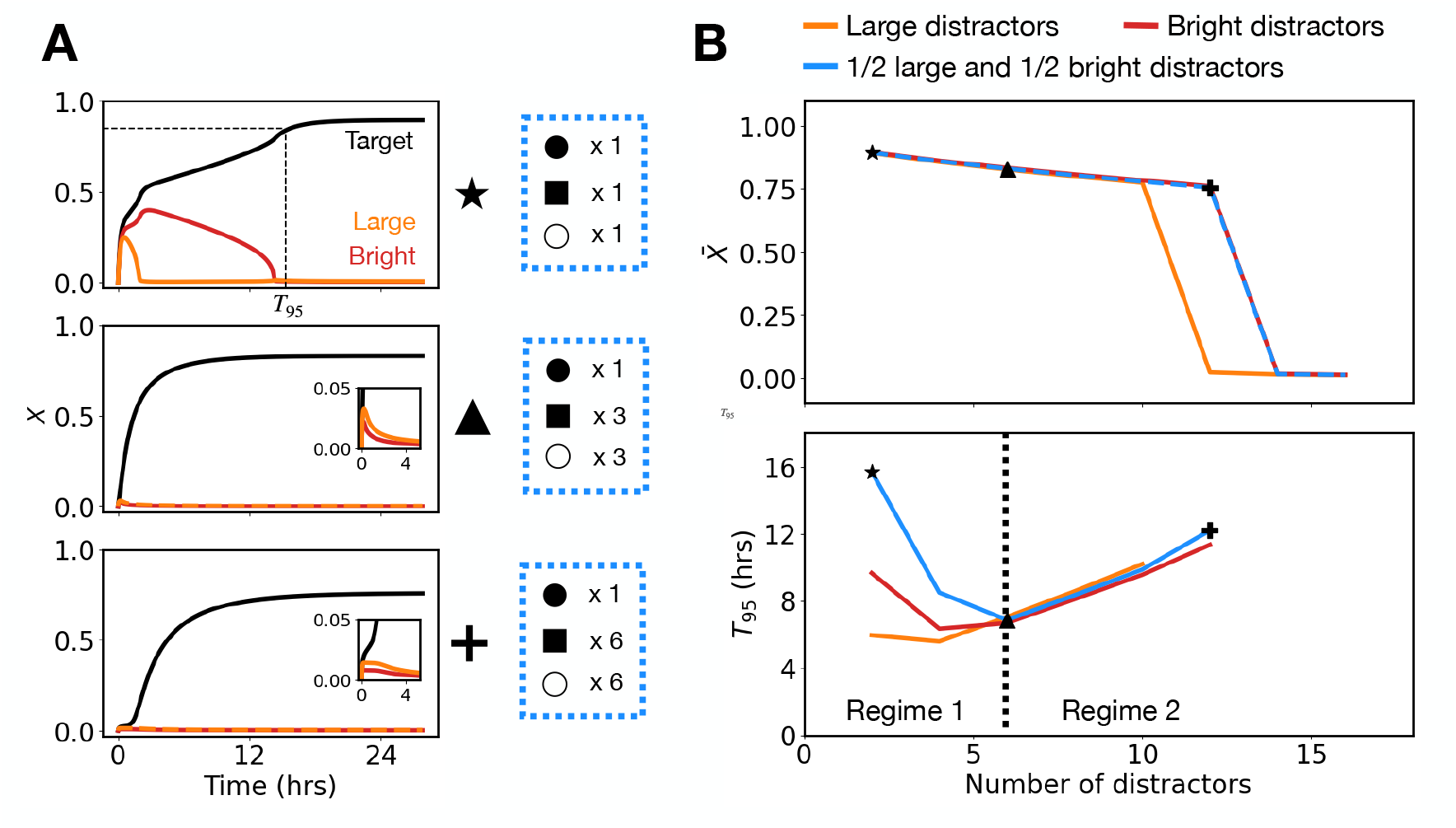
Dynamics and properties of the mean-field model. A) Dynamics of the proportion of the group under each shelter (*X*) with 2, 6, and 12 distractors (from top to bottom) in the half-large, half-bright distractor configuration. Black circles represent the target shelter. Black squares represent large distractors. White circles represent bright distractors. B) Proportion of the group under the target shelter after the system converges to equilibrium. (top; 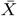) for different distractor configurations. Decision time, defined as the time taken to reach 95% completion (bottom; *T*_95_). Dashed black line represents critical *N* value between the two processing regimes.

Firstly, regardless of distractor types, the model predicts that cockroach groups are able to successfully identify the target shelter at low distractor numbers (Fig. 3B top panel). However, beyond a threshold number of distractors, group consensus breaks down and the final proportion under the target shelter is no more than if cockroaches randomly distributed among all shelters, with the location of this threshold dependent on the distractor type. We refer to this threshold as the collapse point.

Secondly, the model predicts that the relationship between the *T*_95_ decision time (the time taken to reach 95% of the final proportion) and the number of distractors is non-monotonic, as already seen in Figure 3A. Figure 3B (bottom panel) shows that at low numbers of distractors (we call this Regime 1), the decision time decreases as distractor number increases. Beyond a critical number of distractors (the value of this depends on the type of distractors present), the decision time increases as the number of distractors increase (we call this Regime 2). This non-monotonic relationship is surprising – one might naively expect a monotonic increase in decision time as the number of distractors increase due to the increase in decision complexity. This therefore suggests the existence of competing processes during information processing that dominate at different levels of decision complexity.

Finally, having a mix of distractors that are unattractive in either the capacity or brightness attribute (1/2 large and 1/2 bright distractors case) is a more complex problem than if all distractors were only unattractive in the same attribute (large distractors or bright distractors case). This is because multiple attributes would need to be integrated to solve the best-of-*N* task (see Fig. 1). Intuitively, we therefore expected that these decisions would take longer and that the group would become unable to make a decision at a lower number of distractors. We found that when distractor types are mixed, the decision time is indeed higher at low numbers of distractors. However, the decision times become very similar across distractor configurations at high distractor numbers. In fact, the decision time of the mixed distractor configuration is marginally lower than the all large distractors configuration when there are 6, 8, and 10 distractors. In addition, when there are mixed distractors, the system is able to support collective decisions up to 12 distractors, higher than that compared to when there are only large distractors and equal to only bright distractors. Thus, it seems that higher decision complexities do not necessarily translate to slower and less effective decision-making.

#### 3.3.2 Explaining the dynamic mechanism underlying the collapse point

When all distractors are of the same type (all large distractors and all bright distractors configuration), the collapse point of the system can be mathematically derived by reducing the mean-field model with low decision complexity configurations to two equations for the variables *X*_*T*_ and *Y* (*X*_*T*_ is the proportion of the group under the target shelter, *Y* is the proportion of the group under all distractor shelters; see A). The corresponding differential equations govern the change in the proportion of the group under the target and all distractor shelters, respectively. This reduction is possible since all distractors of the same type follow identical dynamics and therefore the variables defining their dynamics can be lumped together into one.

Examining the phase portraits in Figure 4A, the shapes of the nullclines as well as the location, number and stability properties of the fixed-points depend on the number of distractors and other model parameters.

**Figure 4:**
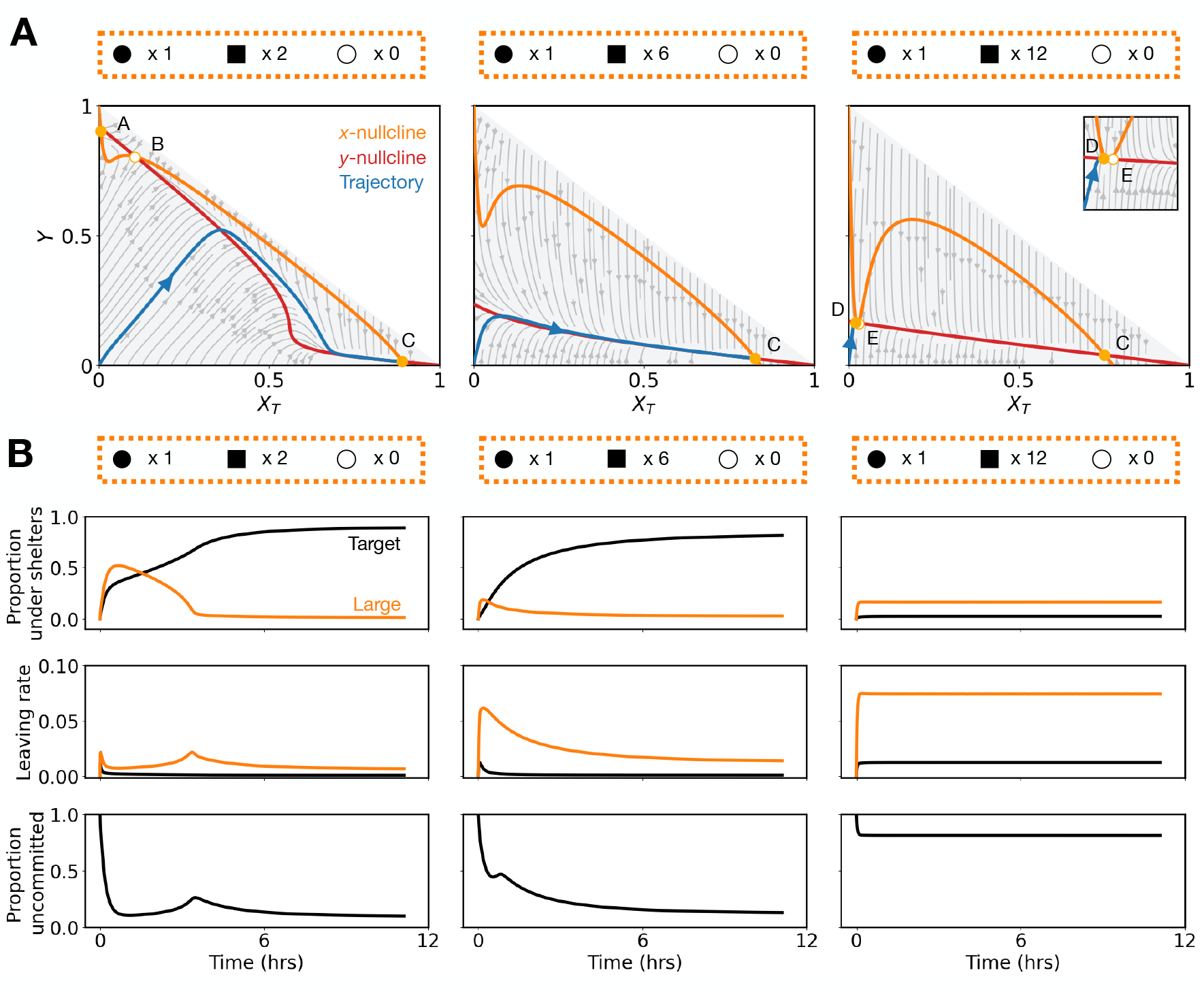
Phase portraits and dynamics at 2 (left column), 6 (middle column), and 12 (right column) distractors in the all large distractors configuration. A) Phase portraits. *X*_*T*_ is the proportion of the group under the target shelter and *Y* is the proportion of the group under all distractor shelters. Green and red lines represent the *X*_*T*_ - and *Y* -nullclines, respectively. Gray arrows represent vector fields and gray curves represent trajectories of the system for different initial conditions. The trajectory when the system starts with all cockroaches uncommitted is shown in blue. Shaded gray area represents feasible regions in the phase plane, since the proportion under the target and the proportion under the distractors needs to sum to 1. Orange dots (labelled A-E) are the fixed points of the system. Stable fixed points are filled and unstable fixed points are empty. B) Dynamics for different parameters. (Top row) Proportion of group under different shelters over time (*X*). (Middle row) Leaving rate from shelters over time 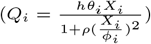. (Bottom row) Number of uncommitted cockroaches over time.

At low numbers of distractors (Figure 4A left panel), there are three fixed-points. Stable fixed points A and C correspond to an even split of the group among the distractors and to a group consensus under the target shelter, respectively. There exists a threshold (separatrix) around the unstable fixed point B that determines which stable fixed point the trajectory evolves towards (depending on the initial condition). The trajectory lies on the right of the threshold, and tends towards C.

As the number of distractors increases, the system undergoes a saddle-node bifurcation, where fixed points A and B collide and are destroyed, leaving only one stable fixed point C (Figure 4A middle panel), towards which the trajectory must converge.

As the number of distractors continues to increase, beyond the collapse point (Figure 4A right panel), the system undergoes another bifurcation where a stable fixed point (D) and an unstable fixed point (E) is created with a low *X*_*T*_ and *Y* . As this happens, the vector field becomes “more vertical”, and it directs the trajectory towards D, thus switching the stationary state to this newly created fixed-point. This explains the abrupt transition of the curves in Fig. 3B. The stable fixed point C still exists, but trajectories from the initial uncommitted state never converge to it.

#### 3.3.3 Explaining the non-monotonic relationship between decision time and number of options

We hypothesize that the decrease in decision time at low distractor numbers stems from a reduced effectiveness in the ability of distractors to retain individuals (we call this Effect 1). Initially, cockroaches explore all shelters until the group is approximately evenly distributed across all shelters. We call this the ‘initial exploration phase’. The proportion of individuals under each distractor therefore peaks at approximately 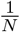. This value decreases as more distractors are added, increasing the leaving rates from the shelters (since the leaving rat 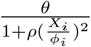 increases with decreasing density). This temporarily enlarges the pool of uncommitted individuals. Since the target shelter retains cockroaches more effectively, the larger uncommitted pool increases recruitment to the target, accelerating consensus. This hypothesis is supported by Fig. 4B. As the number of distractors increases, there is a greater efflux of individuals from the distractors, thereby increasing the size of the uncommitted pool. This is a novel description of a collective decision-making mechanism where increasing decision complexity—up to a point—accelerates consensus.

As the number of distractors continues to increase, however, the likelihood of encountering the target shelter decreases, and recruitment to all shelters is split (we call this Effect 2). In Regime 2, Effect 2 dominates over Effect 1 and slows consensus formation under the target shelter.

As the leaving rate is inversely proportional to the square of the density term, the strength of Effect 1 (weaker distractors) on reducing the decision time is proportional to 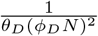. Meanwhile, the strength of Effect 2 (reduced probability of finding a target) on increasing decision time is proportional to 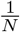. Since the strength of Effect 1 decays faster with *N*, Effect 2 dominates at higher distractor numbers. The fact that the strength of Effects 1 and 2 depend on *N* and scale differently explains why two processing regimes exist.

#### 3.3.4 Explaining the two distractor types case

Examining the dynamics when there are two distractor types reveals another novel emergent filtering mechanism. Here, we use the term “filtering” to describe the proportion of the group under a distractor shelter decreasing to zero. The rate of filtering depends on the attributes of the distractor and the number of distractors. This is seen in Fig. 3A, which shows that after the initial exploration phase, the proportion of the group under a distractor type ultimately decreases to 0 at different rates depending on the distractor type. We will refer to the distractor type that gets filtered faster as a “weak distractor”, and the distractor type that gets filtered slower as a “strong distractor”. While having distractors that differ along two attributes is a higher decision complexity problem, the cockroach group can effectively reduce the decision complexity by filtering the weak distractor first, then selecting the target from the remaining smaller number of strong distractors. At low distractor numbers (under 6 distractors), this process of reducing decision complexity increases the decision time, since in Regime 1, decision time decreases with an increasing number of distractors. At high distractor numbers, this process results in decision times that are higher than if all distractors were of the weak type, and lower than if all distractors were of the strong type, since the weak distractors get filtered faster than the strong distractors.

To test this, we systematically changed the ratio of large to bright distractors (Figure 5A) and compared their *T*_95_ decision times. The results show that whenever there is a mixture of distractor types (when the ratio of large:bright distractors is 3:1, 2:2, or 1:3) at low and high numbers of distractors, there is a gradual decrease in *T*_95_ decision times as the ratio of large to bright distractors decreases, consistent with our hypothesis.

**Figure 5:**
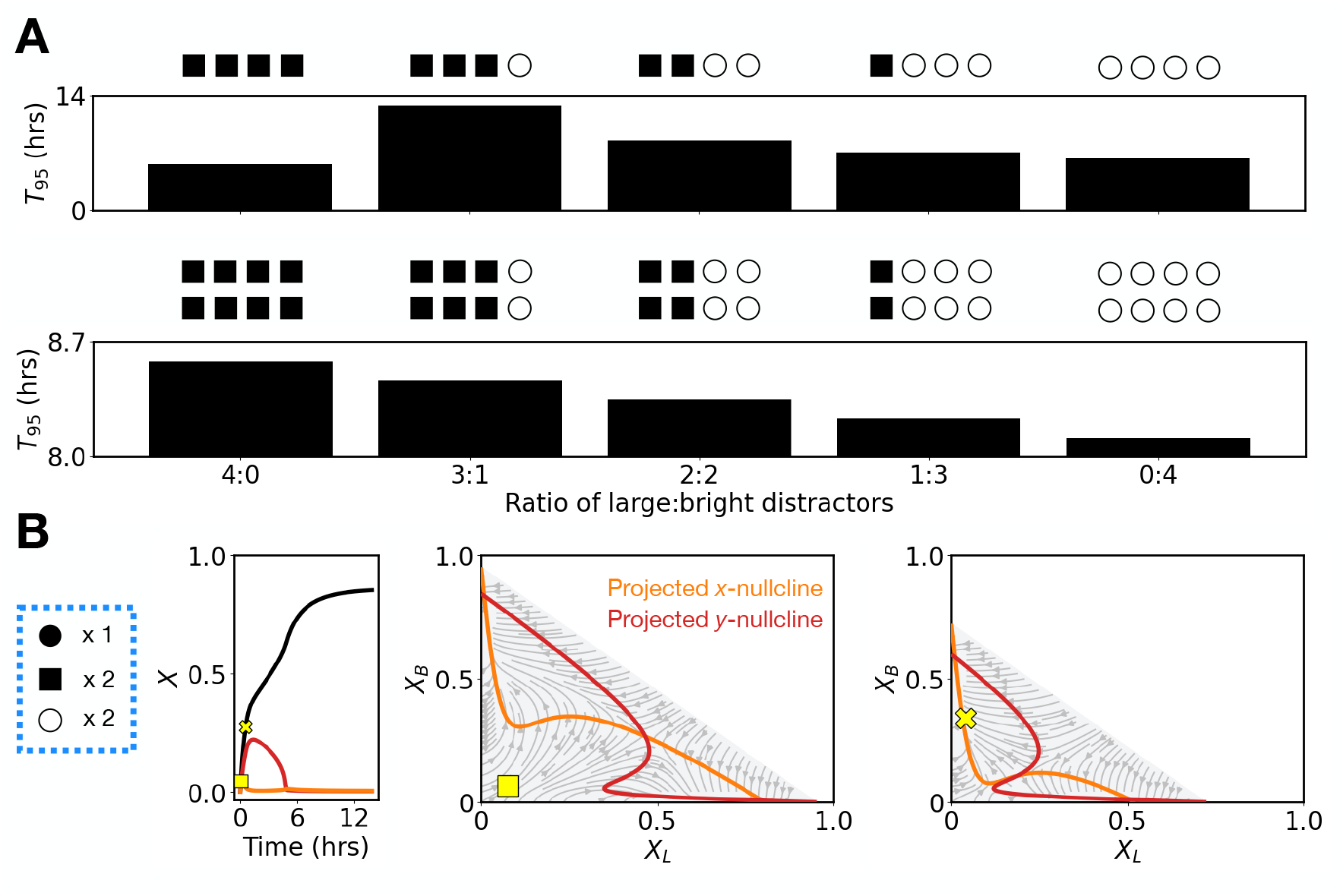
A) Decision time at different ratios of large to bright distractors. Simulations were performed with different ratios of large distractors (denoted by black squares) and bright distractors (denoted by white circles). The ratios tested were 4:0, 3:1, 2:2, 1:3, and 0:4 with 4 total distractors (top panel) and 8 total distractors (bottom panel). B) Proportion of the group under each shelter with 6 distractors for the half large and half bright distractor configuration and the corresponding phase portraits with the proportion under the target fixed at two values corresponding to the yellow square and yellow cross markers. *X*_*L*_ and *X*_*B*_ are the proportions of the group under all the large and all the bright distractors respectively. Green and red lines represent the projected *x* - and *y* -nullclines respectively. Shaded gray area represents feasible regions in the phase plane, since the proportion under the target and the proportion under the distractors needs to sum to 1.

We can understand this mathematically by reducing our system to 3 equations by describing the population of shelters with the same attributes with a single equation (see B). Thus, we have three equations governing the evolution of *X*_*T*_ (proportion of the group under the target shelter), *X*_*L*_ (proportion of the group under all the large distractors), and *X*_*B*_ (proportion of the group under all the bright distractors).

The phase portraits for this system are three-dimensional, which is difficult to visualize. To aid in the analysis, we slice the phase portraits along constant values of *X*_*T*_ and we track the evolution of these projections (of the 3D space onto the *X*_*T*_ -planes) as *X*_*T*_ monotonically increases (Fig. 5B left).

Figure 5B shows such projections for two representative values of *X*_*T*_ where the yellow square (middle) and yellow cross (right) indicate the state of the system at the corresponding time (left).

The first value of *X*_*T*_ (yellow square) represents the initial exploration phase, so the group is distributing itself roughly equally between all shelters. Consistent with this, around the yellow square, the vector field points away from zero. The vector field directions are unequally distributed (while it is biased upwards, it implies that a small perturbation to *X*_*L*_ at this point in time will change the trajectory towards the right). The yellow square is mounted on a vector field pointing upwards. This causes the trajectory to move towards the higher values of *X*_*B*_ and lower values of *X*_*L*_. This is clearly seen for the value of *X*_*T*_ corresponding to the yellow cross, which shows that the group has eliminated large distractors from the competition. Beyond these two points, the region of the phase portraits where the projected nullclines are located increasingly shrinks, as the trajectory of interest reaches its maximum value of *X*_*B*_ and the proportion of the group under both distractor types decreases to zero (not shown).

The specific directionality in the vector fields shows that cockroach groups are utilizing differential filtering rates to reduce the decision complexity of the search problem.

### 3.4 Sensitivity analysis predicts a singular processing regime when distractors are easier

In our results above with a single distractor type, we have been simulating best-of-*N* tasks with either *ϕ*_*D*_ = 1.75 or *θ*_*D*_ = 1.75 (i.e., either the distractors are all 1.75 less attractive than the target in the capacity attribute or in the brightness attribute). Do the qualitative results obtained still hold with different values of *ϕ*_*D*_ and *θ*_*D*_?

Since the strength of Effect 1 (described above) is inversely proportional to *ϕ*_*D*_ and *θ*_*D*_, we predicted that if either *ϕ*_*D*_ or *θ*_*D*_ is large (i.e., there is more contrast between the target and distractor shelter), Effect 1 should be weak even at low distractor numbers and Effect 2 should dominate. This implies that we should only observe increasing decision times with distractor number. We tested this with a sensitivity analysis.

In Fig. 6, we consider the case where all distractors are unattractive in the capacity attribute (*ϕ >* 1). We measured how the final proportion and *T*_95_ decision times scale with distractor numbers at low and high values of *ϕ*_*D*_ and found that at high *ϕ*_*D*_, decision time increases monotonically with distractor number (Fig. 6A). This confirms that Effect 1 has been dominated by Effect 2. To quantify this behavior systematically, we performed a sweep of 50 different values of *ϕ*_*D*_ and measured the number of distractors required to reach the collapse point (*N*_collapse_), the number of distractors where decision time is decreasing (i.e., the size of regime 1; | Regime_1_ |), and the gradient of the increase in decision time in regime 2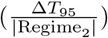.

**Figure 6:**
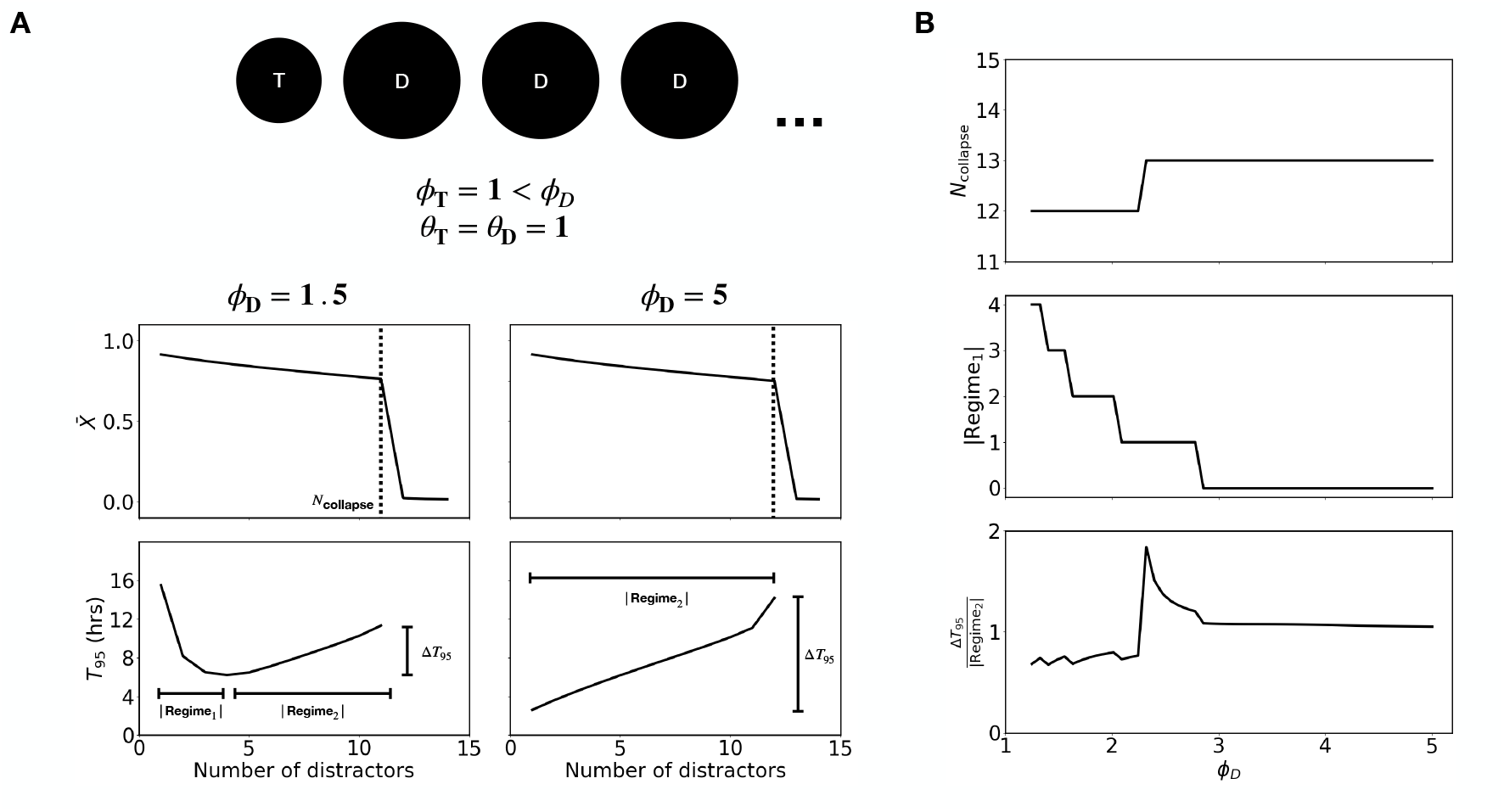
A) (Top) Setup of the sensitivity analysis. Distractors are all unattractive in the capacity attribute, and are *ϕ*_*D*_ times larger than the target shelter. (Bottom) Proportion of the group under the target shelter 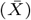 and decision time (*T*_95_) graphs for *ϕ*_*D*_ = 1.5 and *ϕ*_*D*_ = 5. The collapse point (*N*_collapse_) is the maximum number of distractors for which the group is still able to aggregate under the target shelter. Regime 1 size is measured as the number of distractors that show decreasing decision times. Regime 2 gradient is calculated by dividing the total difference in decision time in Regime 2 by the size of Regime 2. B) Plots showing collapse point (top), Regime 1 size (middle) and Regime 2 gradient (bottom) for 50 values of *ϕ*_*D*_ from 1.25 to 5.

These values are plotted in Fig. 6B. We found that the collapse point (top) only slightly increases with *ϕ*_*D*_. From the phase portraits, increasing *ϕ*_*D*_ changes the vector field by moving the nullclines (Supplementary Figure 2). The saddle-node bifurcation that underlies the collapse point now occurs at a slightly higher distractor number.

As expected, we found that the size of Regime 1 decreases gradually with *ϕ*_*D*_. Intuitively, the weakening of distractor strength in Effect 1 only works if distractors are individually strong enough to compete with the target: weakening a distractor is not a significant influence on decision time if it is already weak to begin with. Thus, the higher the *ϕ*_*D*_, the weaker the distractor, and the less significant Effect 1 is compared to Effect 2.

The gradient of Regime 2 (bottom) is non-monotonic in *ϕ*_*D*_ and does not change drastically, remaining close to 1 across *ϕ*_*D*_ values. This is because Effect 2 scales with 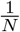, so it does not depend on *ϕ*_*D*_. The spike in gradient is due to the increase in the position of the collapse point. At the higher collapse point, it takes longer for the target shelter to reach the threshold where positive feedback takes effect. This results in a sharp increase in decision time and thereby in the Regime 2 gradient.

We performed the same sensitivity analysis for the case where all distractors are unattractive in the brightness attribute (*θ*_*D*_ *>* 1) and showed the same qualitative results (Supplementary Figure 3).

### 3.5 Agent-based model confirms mean-field model predictions

From Fig. 7A, the agent-based model (ABM) simulations quantitatively match the mean-field model. We can also examine the dynamics of shelter elimination by measuring the order of elimination for large compared to bright distractors. The elimination order is measured by recording the order in which the number of individuals under each shelter reaches its peak in a simulation.

**Figure 7:**
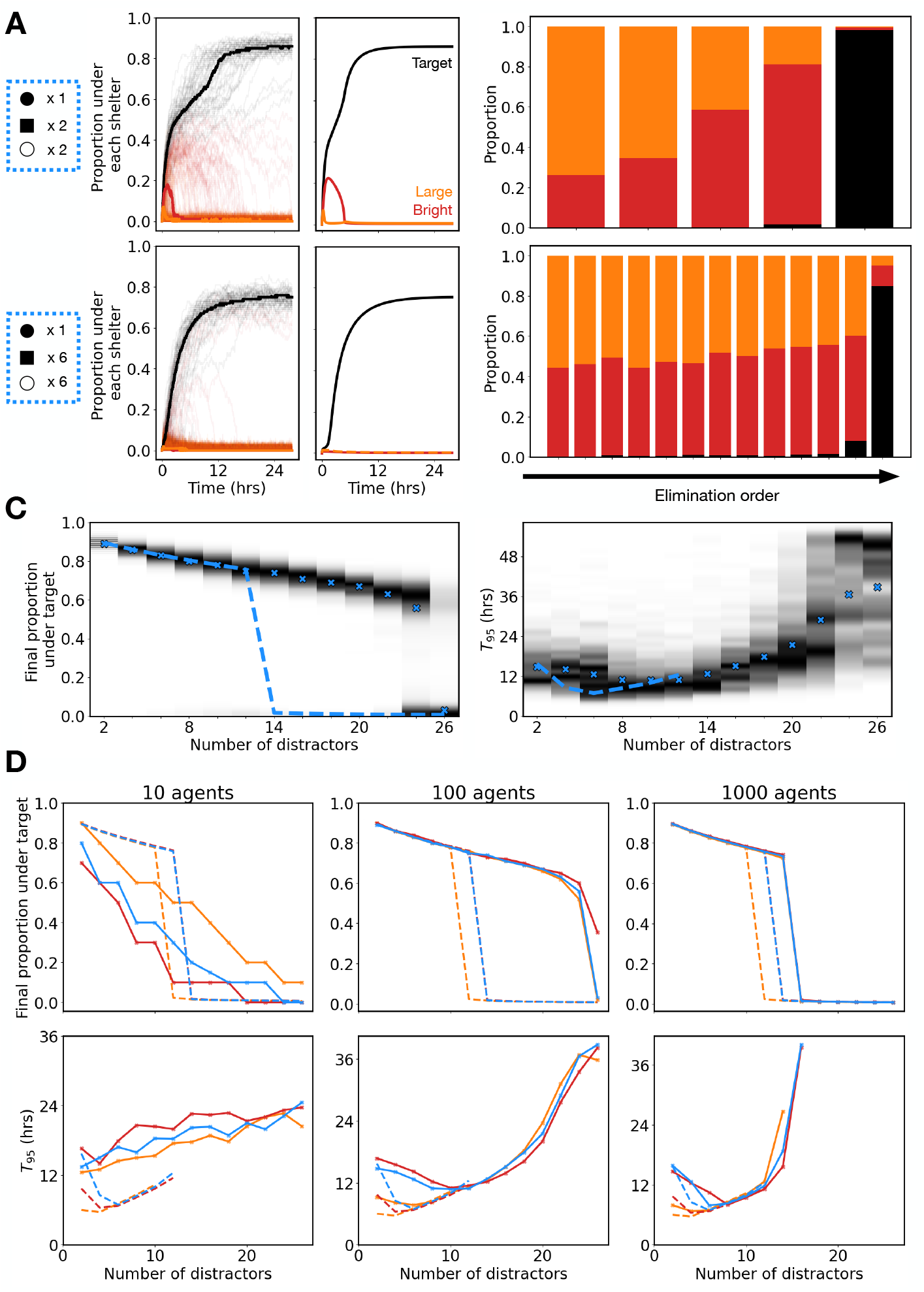
A) (left panel) ABM simulations with 4 and 12 distractors for the half large and half bright distractor configuration compared to mean-field model simulations. Solid lines represents the median over 1000 simulations. 50 randomly chosen raw simulations are shown in the background with low transparency. (right panel) Shelter elimination dynamics averaged over 100 simulations. Each stacked bar shows proportion of distractor types eliminated in that position. C) Distribution of the final proportion under the target shelter and *T*_95_ decision times for the half large and half bright distractor configuration across different numbers of distractors. The distributions are taken by simulating the same model at each number of distractors 500 times. Dark areas represent peaks in the distribution and crosses represent the median. Dotted line shows the mean-field model predictions. D) Summary of final proportion under the target shelter and *T*_95_ decision times across different distractor configurations and different numbers of agents.

The earlier a shelter reaches its peak, the earlier it decays and is therefore “eliminated” earlier. From Fig. 7A (right panel), large distractors are typically eliminated first before bright distractors at lower numbers of distractors. At higher numbers of distractors, this pattern disappears and there is an equal chance of either distractor type getting eliminated first.

The reason for this is that at the beginning of simulations, agents will tend to distribute themselves randomly across all shelters. At low distractor numbers, each shelter is more populated, meaning that each distractor is more competitive. Elimination of each distractor therefore strongly depends on shelter quality. At higher distractor numbers, each shelter is barely populated, so elimination of distractors is largely based on random chance.

Figure 7C demonstrates that the ABM successfully replicates the key qualitative behavior observed in the mean-field model. Specifically, decision time exhibits a characteristic non-monotonic relationship with distractor number: it decreases at low distractor numbers and increases beyond a critical value of *N*.

Notably, the ABM has a higher collapse point compared to the mean-field model at higher distractor numbers, being able to converge on the target even with up to 22 distractors, compared to only 12 distractors in the mean-field model. While this may seem surprising initially, examining the phase portrait in Fig. 4A shows that at high numbers of distractors, the basin of attraction for the stable fixed point representing indecision is small, meaning that small perturbations to the original uncommitted state are sufficient to push the system to converge towards the stable fixed point representing consensus under the target shelter. The stochasticity inherent in ABMs therefore allow the group to bypass the indecision fixed point and achieve consensus up to much higher distractor numbers.

To test how this changes with group size, we repeated the simulations using 10, 100, and 1000 agents (Fig. 7D). With small group sizes, we observe qualitatively different group-level properties: the final proportion decreases smoothly and decision time increases. This suggests that the mechanisms for the collapse point and decision time decreasing (as described above) requires some minimum number of cockroaches to take effect. As group size increases, the ABM’s behavior smoothly converges to that of the mean-field model, both in terms of the proportion under the target shelter and in decision time. This is consistent with expectations – since the mean-field model is a large-scale approximation of the agent-based system, it becomes increasingly accurate in the large population limit.

### 3.6 Spatial ABM shows qualitatively similar results

In the real world, the spatial arrangement of the shelters may affect the dynamics of collective decision-making as it influences the likelihood of a cockroach discovering a shelter. Whether a shelter is likely to be explored by a cockroach is determined in large part by where the cockroach is – a cockroach will much more likely enter a shelter adjacent to the shelter it exited from compared to a distant shelter. The ABM implemented thus far does not take this into account, since all shelters have equal discovery probabilities.

To test whether including the spatial aspect of transitions qualitatively affects our results, we modify the ABM to include different transition probabilities between shelters. To derive these probabilities, we simulated random walks through a two-dimensional space where shelters are arranged in a triangular lattice (see Fig. 8A left). The transition probabilities were then determined by measuring the frequency of transitions between every pair of shelters (Fig. 8A right).

**Figure 8:**
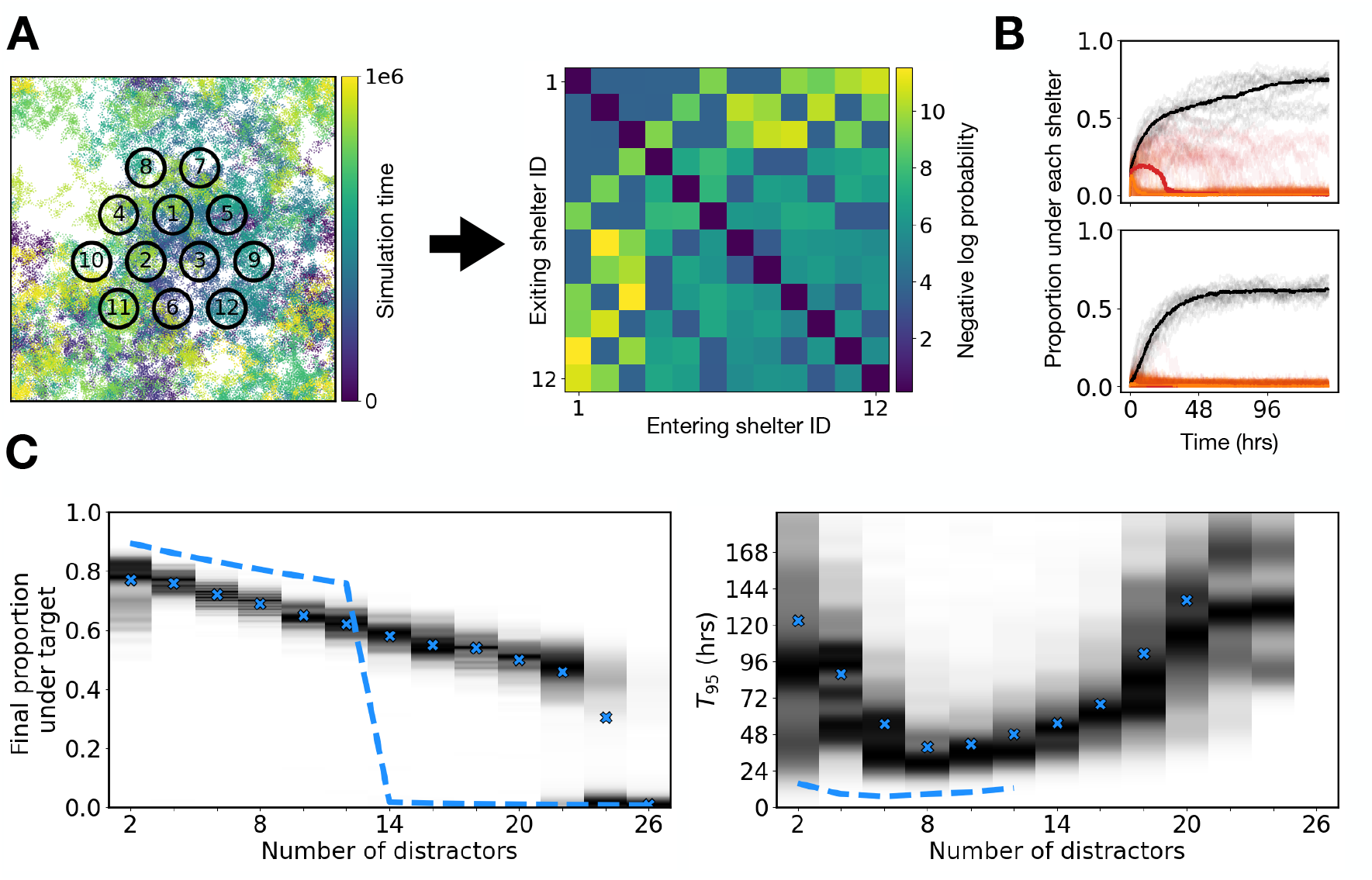
A) Schematic showing derivation of transition matrix. *N* shelters are placed in a triangular grid within a field and a random walk is simulated (left; *N* = 12 is shown). The frequencies of transitioning between every pair of shelter is recorded and transformed into a *N × N* transition matrix. B) Proportion under each shelter with 4 (top) and 12 (bottom) distractors for the half large and half bright distractor configuration. Solid lines represent the median over 200 simulations. 20 randomly chosen raw simulations are shown in the background with low transparency. C) Distribution of the final proportion under the target shelter and *T*_95_ decision times for the half large and half bright distractor configuration across different numbers of distractors. The distributions are taken by simulating the same model at each number of distractors 500 times. Dark areas represent peaks in the distribution and crosses represent the median. Dotted line shows the mean-field model predictions.

From Fig. 8B, the decision-making dynamics are qualitatively similar to that of the mean-field model and ABM in Fig. 7A. As the number of distractors increase, the group is still able to find the target, with the final proportion under the target shelter following a similar decreasing trend (Fig. 8C left panel). Quantitatively, the decision times of the spatial ABM are around doubled compared to the original ABM, which is intuitive as decreasing the likelihood of discovering other shelters will inevitably delay the decision-making process. Importantly, the decision times show a similar decrease in decision time at low to intermediate numbers of distractors, and then an increase as the distractor number continues to increase, showing that the addition of a spatial component to the model does not qualitatively change the results shown from the other models.

## 4 Discussion

Collective decision-making experiments in cockroach groups have largely focused on simple binary choices with a single differing attribute (Amé et al., 2006; Canonge et al., 2009). Here, we have addressed the question of how decision-making dynamics scale with decision complexity by analyzing a classical dynamical model of shelter selection in cockroach groups. By examining how the complexity of the task influences group-level decision-making properties like decision accuracy and decision time, we show a number of counterintuitive predictions that cannot be made simply by generalizing results from existing work with best-of-2 experiments.

### Prediction 1: Cockroaches value shelter capacity over shelter brightness

Our model predicts that when comparing shelters that are either unattractive in the capacity attribute or equally unattractive in the brightness attribute, cockroaches prefer the latter. This means that they would prefer to compromise on shelter brightness than shelter capacity. While best-of-2 experiments in cockroaches have been performed with either capacity (Amé et al., 2006) or brightness (Canonge et al., 2009) as a differing attribute, no experiments have been carried out to compare trade-offs and preferences between both attributes.

### Prediction 2: Cockroaches use a compensatory algorithm when processing attribute information

The ability to perform sophisticated trade-off reasoning has long been considered a hallmark of advanced cognition. We showed that cockroaches are willing to compromise on a lower attractiveness in the capacity attribute if the shelter is compensated by a sufficiently attractive brightness attribute. Therefore, we predict that cockroach groups are capable of performing trade-off reasoning, a collective ability that had previously only been shown in house-hunting *Temnothorax* ants (Franks et al., 2003).

Identifying whether animal groups use compensatory or non-compensatory algorithms to make decisions reveals whether trade-off reasoning can emerge from simple social interactions. In house-hunting *Temnothorax* ants, the ability to perform trade-off reasoning at the colony level is simply a result of the fact that individual ants translate an overall assessment of an option into a single ‘score’ that modulates how long it waits before initiating recruitment (they hesitate for longer before recruiting nestmates when they encounter less attractive nest site) (Mallon et al., 2001; Franks et al., 2003). This is the same fundamental principle underlying trade-off reasoning in our model of cockroach shelter selection. Both density 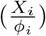 and brightness (*θ*_*i*_) terms are scaling factors that modulate the leaving rate, which directly affects which shelter is selected. A low level of attractiveness in the brightness attribute will increase the leaving rate, but this can be reversed by a high level of attractiveness in the capacity attribute. Intuitively, trade-off reasoning will emerge in any collective system where the values of different option attributes regulate a single individual-level behavior that directly influences consensus formation.

### Prediction 3: Beyond a certain number of distractors, there exists a collapse point where the group becomes unable to decide

Choice overload is a well-studied cognitive phenomenon whereby an increase in the number of available options degrades decision-making in individuals (Schwartz, 2016; Chernev et al., 2015; Tanner and Hemingway, 2025). Whether collective systems like animal groups experience choice overload remains unknown. A theoretical paper modeling honeybees performing symmetric nest-site selection (selecting a nest site among options that all have the same attributes) postulated that honeybees may experience choice overload since the level of signaling necessary to break deadlocks theoretically grows quadratically with the number of options (Reina et al., 2017).

Our model predicts that choice overload also occurs in cockroach groups. Specifically, decision-making breaks down abruptly beyond 12 to 14 distractors as a result of a saddle-node bifurcation in the group dynamics. The bifurcation creates a stable and an unstable fixed point near the initial conditions (all cockroaches uncommitted), preventing the trajectory from entering the basin of attraction for the stable fixed point representing target consensus. This is a novel prediction that, to our knowledge, has not been experimentally observed in any collective system. It supports the hypothesis that, just like with individuals (Tanner and Hemingway, 2025), there can be cognitive processing limits at the level of the group.

### Prediction 4: The relationship between the number of distractors and decision time is non-monotonic

Examining how decision time changes with the number of distractors revealed two distinct processing regimes. At low distractor numbers, the *T*_95_ decision time decreased as more distractors were added (Regime 1). Beyond a certain distractor threshold (we show this to be 4 to 6 depending on the type of distractors), the decision time increases with distractor number (Regime 2).

This phenomenon is due to two underlying effects that influence the decision time in opposite ways as the number of distractors increases. (Effect 1) Every additional distractor competes with other shelters by ‘recruiting’ from the same pool of uncommitted individuals. Since the effectiveness of a shelter in retaining individuals is dependent on the density of individuals under it, adding shelter competition reduces the competitiveness of all distractors, thereby increasing the pool of uncommitted individuals. The target shelter has a lower leaving rate compared to the distractors, so a larger uncommitted pool results in a relatively faster increase in the accumulation of individuals under the target shelter, accelerating consensus. (Effect 2) Every additional shelter decreases the probability of cockroaches finding the target shelter while exploring the environment, reducing the initial rate at which cockroaches enter the target, and thereby increasing the decision time.

These effects scale differently with the number of distractors. The strength of Effect 1 in decreasing decision time is proportional to 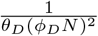, while the strength of Effect 2 in increasing decision time is proportional to 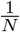. Thus, while Effect 1 dominates at low distractor numbers, it decays faster than Effect 2 with *N*, so Effect 2 dominates at high distractor numbers. This scaling also implies the counterintuitive prediction that decreasing the difficulty of the best-of-*N* problem by increasing the contrast between the target and the distractors (increasing *ϕ*_*D*_ or *θ*_*D*_), will result in an increasingly smaller Regime 1, until only Regime 2 exists.

Collective best-of-*N* search problems are, arguably, conceptually similar to visual search tasks in cognitive psychology. In these tasks, human subjects search for a target object among many distractors that may differ in one (feature search) or several (conjunction search) attributes. In feature search, increasing the number of distractors does not change the decision time since the target object will always ‘pop’ out compared to the distractors; in conjunction search, increasing the number of distractors increases the decision time (Treisman and Gelade, 1980; Treisman, 1977), since it takes time to attend to each distractor and bind its attributes into a single representation. Classical theories of visual selective attention argue that this implies that the human brain can process a single attribute across options in parallel, but multiple attributes need to be processed in series using attention (Treisman et al., 1977; Treisman and Gelade, 1980; Treisman, 1994).

Although originally developed to explain individual perceptual processes, this framework offers a useful lens for interpreting collective decision-making in groups facing best-of-*N* search problems. By examining how decision time scales with decision complexity, it is possible to infer the extent to which animal groups compare options in parallel. Our model predicts that at low numbers of distractors, cockroach groups process in a parallel-like regime, and transition to a serial regime at high numbers of distractors. This non-monotonic relationship exists because of the nature of local interactions employed by cockroaches. Instead of using coordinated search strategies that require the group to remain as a cohesive unit (like sequential binary search (Sridhar et al., 2021)), cockroaches, like other social insect colonies, can search and evaluate many options simultaneously. In addition, cockroaches do not broadcast any positive feedback signals (like pheromones) to recruit individuals to shelters. Rather, their presence in a shelter acts as a social signal for others to remain in the same shelter (Jeanson et al., 2005). This specific local interaction underlies Effect 1, because fewer individuals exploring a distractor is a direct weakening of that distractor’s ability to compete with the target shelter.

### Prediction 5: When distractors differ across multiple attributes, the group is able to reduce the decision complexity by filtering out weak distractors first

Integrating the different attributes of several options and comparing them to find the best one is a well-studied problem in decision theory, operations research, and cognitive psychology (Triantaphyllou, 2000; Feng et al., 2022; Azimi and Chen, 2025; Payne et al., 2012; Franks et al., 2003). In Prediction 2, we showed that the first aspect—integration of different attributes—is accomplished using a compensatory algorithm that weights different attributes by modulating the leaving rates. In Prediction 4, we showed that the second aspect—comparing options—is accomplished through simultaneous exploration of options and filtering of distractors. Together, these properties imply that cockroach groups solve best-of-*N* problems involving both capacity and brightness attributes in fundamentally the same way as if only one attribute were involved: by eliminating shelters that have relatively high leaving rates. Since there are two types of distractors, the weaker distractor type gets filtered faster, allowing the group to effectively reduce the decision complexity and find the target among a smaller number of distractors. From Prediction 4, reducing decision complexity increases decision time at low distractor numbers and decreases decision time at high distractor numbers. The model therefore predicts that with two distractor types at low distractor numbers, decision time is higher than with one distractor type, and intermediate at high distractor numbers.

In summary, by exploring higher levels of decision complexity, we have shown that our model exhibits sophisticated group-level properties beyond symmetry-breaking between two options. The predictions made by our model not only show that studying best-of-2 experiments alone is insufficient to understand collective decision-making dynamics, but more importantly, that exploring higher decision complexity problems is *necessary* to infer group-level information processing capabilities. This raises the question: if we apply this framework to other models of collective decision-making (e.g., models of mass recruitment in Argentine ants (Goss et al., 1989; Beckers et al., 1990), waggle dancing in honeybees Seeley et al. (2012); Reina et al. (2017), opinion polarization in humans), would we observe similar displays of rich group-level behavior?

## A One distractor type

Since all distractors are of the same type, there are only two distinct dynamics: the target shelter, and the distractors. Let *X*_*T*_ and *X*_*D*_ be the number of individuals under the target and distractor shelters respectively. From equation 1, the target shelter obeys

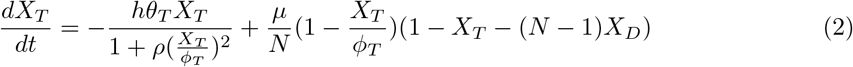

Similarly, the distractors obey

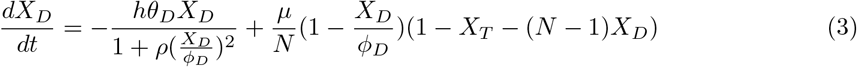

Let 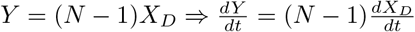

Thus, equations 2 and 3 become

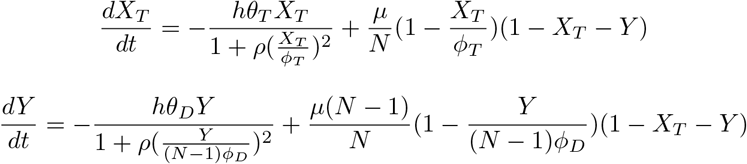

Equating to 0 and solving to find the nullclines

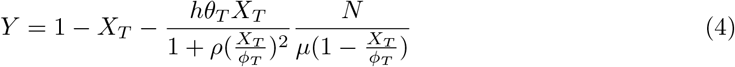

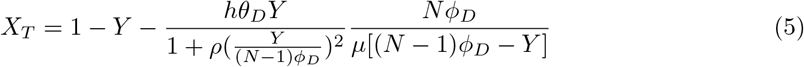

## B Two distractor types

Since there are two distractor types, there are three distinct dynamics: the target shelter, the large distractors, and the bright distractors. Let *X*_*L*_ and *X*_*B*_ be the number of individuals under the large and bright shelters respectively.

From equation 1, the target shelter obeys

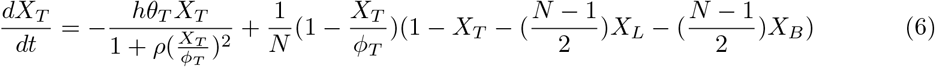

Similarly, the distractors obey

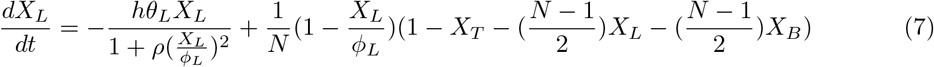

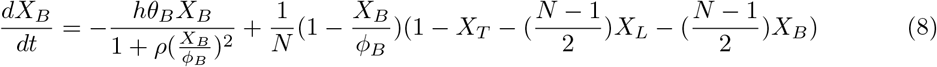

Let 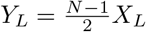 and 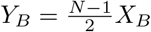

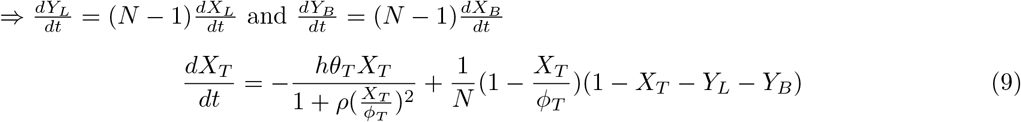

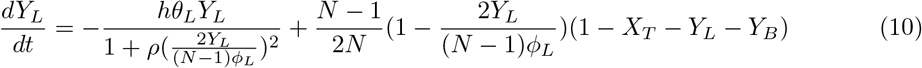

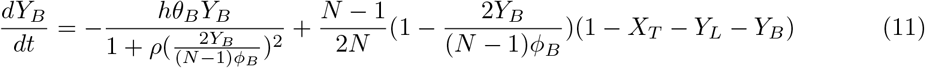

