## Supplementary Materials for "How do Cockroach Groups Integrate Multiple Attributes in a Best-of-N Task?"

### Supplementary Information

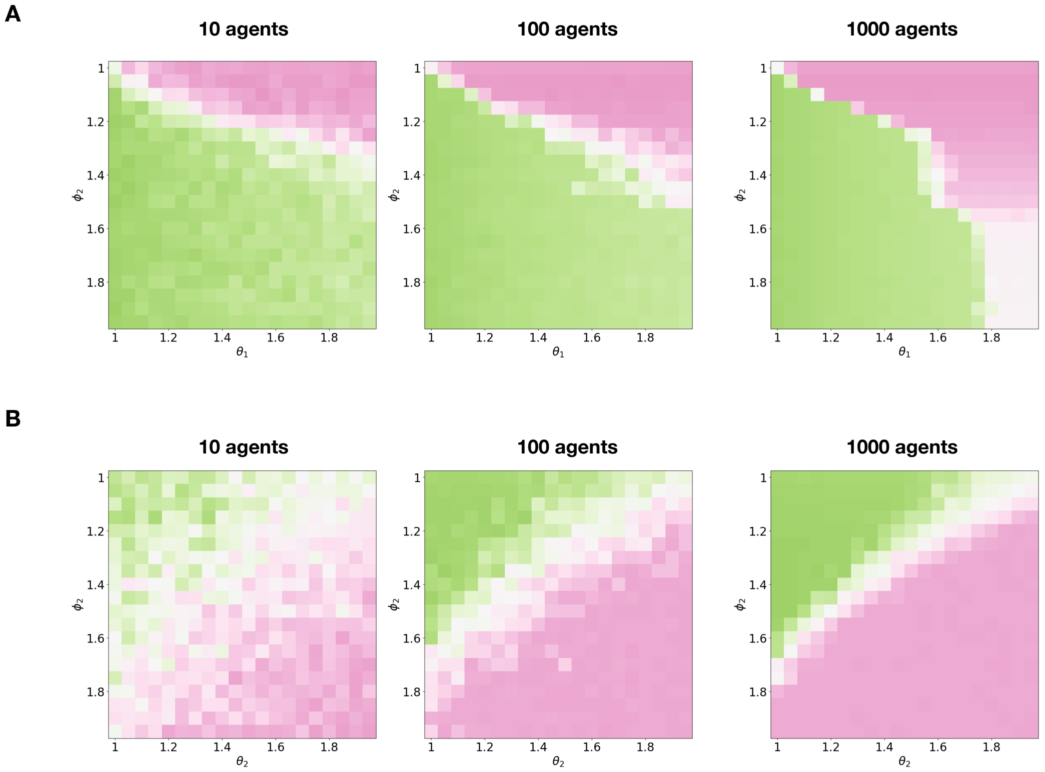


Figure 1: Heatmaps showing which shelter is picked by the agent-based model (ABM) for corresponding experiments to the mean-field model. The ABM model results converge to the mean-field model results as the number of agents increases. A) Experiment 1. Green represents a shelter that is attractive in the capacity attribute ($\phi_1 = 1$) but not in the brightness attribute ($\theta_1 > 1$). Pink represents a shelter that is attractive in the brightness attribute ($\theta_2 = 1$) but not in the capacity attribute ($\phi_2 > 1$). B) Experiment 2. Green represents a shelter that is attractive in the capacity attribute ($\phi_1 = 1$) but not in the brightness attribute ($\theta_1 = 2$). Pink represents a shelter that is less attractive in the capacity attribute ($\phi_2 > 1$) but more attractive in the brightness attribute ($1 < \theta_2 < 2$).

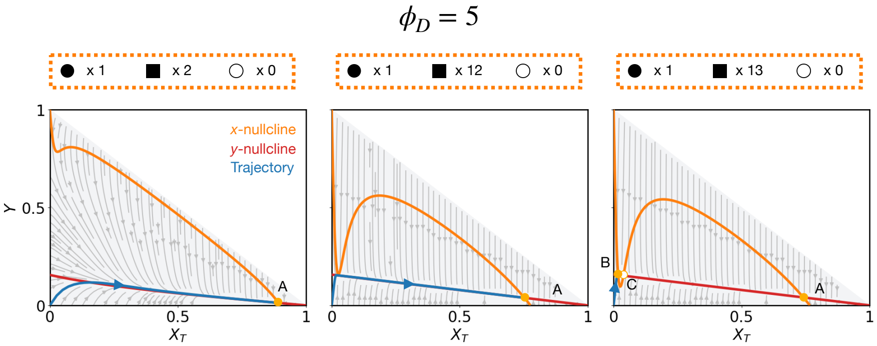


Figure 2: Phase portraits for all large distractors case with $\phi_D = 5$ at 2 (left), 12 (middle), and 13 (right) distractors. $X_T$ is the proportion of the group under the target shelter and $Y$ is the proportion of the group under all distractor shelters. Green and red lines represent the $X_T$- and $Y$-nullclines, respectively. Gray arrows represent vector fields and gray curves represent trajectories of the system for different initial conditions. The trajectory when the system starts with all cockroaches uncommitted is shown in blue. Shaded gray area represents feasible regions in the phase plane, since the proportion under the target and the proportion under the distractors needs to sum to 1. Orange dots (labelled A-C) are the fixed points of the system. Stable fixed points are filled and unstable fixed points are empty.

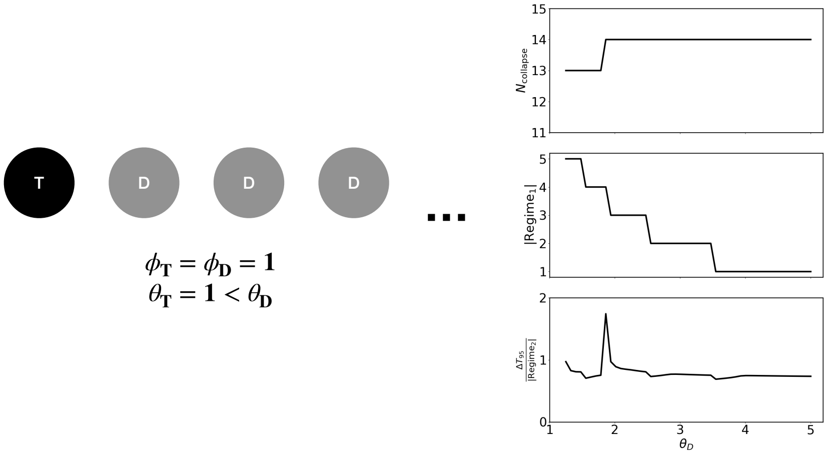


Figure 3: (Left) Setup of the sensitivity analysis. Distractors are all unattractive in the capacity attribute, and are $\phi_D$ times larger than the target shelter. (Right) Plots showing collapse point (top), Regime 1 size (middle) and Regime 2 gradient (bottom) for 50 values of $\phi_D$ from 1.25 to 5.
